# Lung Disease Network Reveals the Impact of Comorbidity on SARS-CoV-2 infection

**DOI:** 10.1101/2020.05.13.092577

**Authors:** Asim Bikas Das

## Abstract

Higher mortality of COVID19 patients with comorbidity is the formidable challenge faced by the health care system. In response to the present crisis, understanding the molecular basis of comorbidity is essential to accelerate the development of potential drugs. To address this, we have measured the genetic association between COVID19 and various lung disorders and observed a remarkable resemblance. 141 lung disorders directly or indirectly linked to COVID19 result in a high-density disease-disease association network that shows a small-world property. The clustering of many lung diseases with COVID19 demonstrates a greater complexity and severity of SARS-CoV-2 infection. Furthermore, our results show that the functional protein-protein interaction modules involved RNA and protein metabolism, substantially hijacked by SARS-CoV-2, are connected to several lung disorders. Therefore we recommend targeting the components of these modules to inhibit the viral growth and improve the clinical conditions in comorbidity.

## Introduction

The novel Coronavirus Disease 2019 (COVID-19) cases crossed the 4200000 all over the world as of May 12, 2020. The recent data show that the most affected groups are with two or more pre-existing medical conditions such as hypertension, diabetes, metabolic, cardiovascular, and digestive disorder[1–3]. Moreover, comorbidity (or existence of multiple disorders) causes a higher risk of developing a severe illness, poor prognosis, and higher mortality of COVID-19 patients [4]. A virus causes the disease by hijacking the host cell machinery for its replication. Interactions of the virus with host perturb the highly organized host cellular networks and re-construct different networks that are favorable to virus replication.

Similarly, coordinated interactions between molecules in a healthy cell are altered in disease state due to changes in the genetic and epigenetic factors. Hence SARS-CoV-2 interaction pattern with healthy human cells will be different from the disease cell, and this could lead to various impacts on SARS-CoV-2 infection. Human diseases are connected via defects in common genes [5,6]. Moreover, the similarity in disease phenotype often indicates underlying genetic connections. Therefore pre-existing medical conditions can facilitate the appearance of another disease if they share the same or functionally related genes [7]. SARS-CoV-2 has been associated with respiratory tract infection (RTI), and in some cases, it severely damages adult lungs. Here, we predict the risk of COVID19 infection in patients with various lung diseases. In the present work, we have considered a disease in the lung or diseases in other tissues or organs affecting lungs as ‘lung disease’. Recent efforts by Gordon et al. [8] identified 26 of the 29 SARS-CoV-2 proteins, which bind to 332 human proteins and hijack the host translational machinery. Here, we have constructed a tissue (lungs)-specific neighborhood network of the 332 human targets of SARS-CoV-2. Based on the shared genes, we have integrated this neighborhood network with lung diseases and constructed a disease-gene network of the lung. Subsequently, we have built a human lung disease network (HLDN), which also includes COVID19. We observed that 141 lung diseases are associated with COVID19. 49 out of 141 disorders are directly linked to COVID19, apparently justifying the characteristics of a complex disorder. Importantly, HLDN represents a small-world like property, indicating a high-density diseases cluster, indicating severe health risk of patients with comorbidity on SARS-CoV-2 infection.

Next, we identified functional protein modules that are maximally perturbed by SARS-CoV-2 and involved in RNA processing, export, and protein synthesis machinery of the cell. Moreover, these protein modules are associated with various lung disorders, indicating the hotspot for comorbidity. Therefore we suggest targeting these functional protein modules to inhibit the viral growth and improve the clinical conditions in comorbidity. The rate of mutation of SARS-CoV-2 is very high, which enables the virus to develop drug resistance [9]. Therefore identification and targeting host factors will be an enduring approach instead of targeting viral proteins.

## Results

### Construction of SARS-CoV-2 –host interactome in lung

To depict the SARS-CoV-2 –host interaction network, the protein-protein interactions (PPI) network of lungs was obtained from the TissueNet v.2 database [10]. We collected the list of 332 human targets of SARS-CoV-2 from Gordon et al. [8] article and constructed the subnetwork of these 332 proteins from the PPI network of lungs. Out of the 332 viral targets, 323 proteins were present in the subnetwork. The resulting subnetwork, named as SARS-CoV-2 target network (STN), consists of 5050 nodes and 11256 pairwise interactions (Fig.1 a, supplementary table1). The degree distribution of STN demonstrated that it has the scale-free property (Fig. 1b). The network was validated by comparing the average path length of STN with 1000 Erdős–Rényi random graphs of the same density. We observed that the average path length distribution of the 1000 random networks was significantly high (p-value<0.0001) than STN (Fig.1c). Next, we computed the dyadicity (D) (a measure of the connectedness of the nodes with the same label, see method) among the SARS-CoV-2 targets in the STN to know if they share more or fewer edges than expected in a random configuration of the network. We found D=7.664, indicating high connectedness among SARS-CoV-2 targets, or they are aggregated in the same network vicinity. D>1 signifies that SARS-CoV-2 targets are forming a community like structure to hijack the host cellular machinery. Proteins in a community, if implicated in diseases, then they can exhibit a higher chance of comorbidity than those who are not in the community. This is because proteins in a community frequently interact, coexpress, and are functionally interconnected [11]. Therefore to understand the risk of COVID19 with comorbidity, we have constructed and analyzed the disease-gene and disease –disease association map of the STN.

**Fig. 1.**
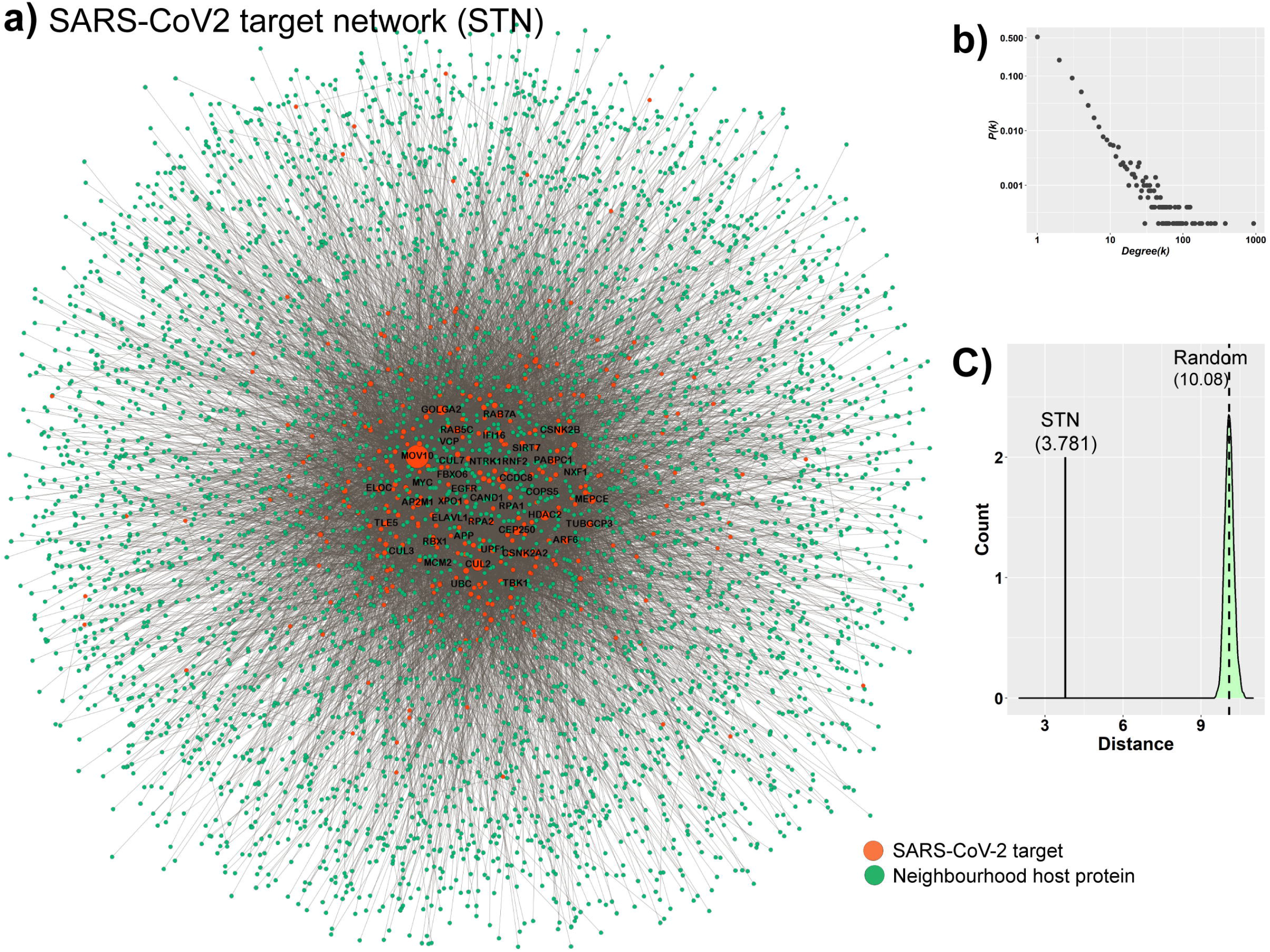
a) Neighbourhood interaction network of SARS-CoV-2 targets (STN) in the lung. The size of the node is proportional to its degree. b) Scatter plot shows the degree distribution of STN. *P(k)* represents the probability of a node with degree *k*. c) The average path length between the nodes in STN and distribution of average path length of 1000 random networks (green), respectively.

### Disease-gene and disease –disease associations map in lungs

To construct a disease association map of STN, we obtained the disease-gene association data from the ORGANizer database [12]. 184 lung diseases, 1957 genes, and 6039 disease-gene pairs were considered for further analysis (see Methods) (supplementary table 2). To construct the disease-gene association map, we screened the diseases which are associated with proteins (nodes) in STN. A disease and node are then connected if the node is associated with the disorder in the lungs. We observed, 618 gene/proteins, consisting of 36 SARS-CoV-2 targets, are linked to a total of 146 disorders, which includes COVID19 (supplementary table 3). Figure. 2a shows the resulting disease-gene association map, named as the lung disease-gene network (LDGN), consisting of 1814 disease-gene pairs. The largest connected component within the LDGN consists of 141 lung diseases and 610 genes, indicating many of the disorders share the common genotype. For example, SARS-CoV-2 targets, FBN1 (degree, *k*=15), FBLN5 (*k* =11), COMT (*k* =9) and neighbourhood nodes, OFD1 (*k* =19), DNAAF2 (*k* =16), DNAAF5 (*k* =16) are linked to multiple disorders (Fig. 2c). Similarly, a disorder in the LDGN is also connected to multiple genes. For instance, ventricular septal defect (*k* =142), respiratory insufficiency (*k* =133), congestive heart failure (*k* =95), apnea (*k* =63) and hypothyroidism (*k* =60) (Fig. 2b and Fig S1). The disease-gene association pattern in LDGN indicates the molecular connection of COVID19 with a wide range of lung disorders. To comprehend the association between COVID19 and lung diseases, a disease-disease association network (DDAN) was constructed, where two diseases were linked if they share one associated gene (Fig 2d). DDAN consists of a total of 141 diseases (nodes) and 1326 links, indicating a higher clustering between diseases. We observed 49 diseases (red nodes) in DDAN, which are directly connected to COVID19 (yellow node) (Fig. 3d). Jaccard similarity coefficient was computed based on the number of common genes to identify the extent of molecular overlapping between lung diseases and COVID19. There are several diseases, like respiratory insufficiency, congestive heart failure, respiratory failure, ventricular septal defect, mitral regurgitation, and hyperthyroidism, which are closely associated with COVID19 (Fig. S2a and b). Thus patients having these disorders probably are more vulnerable for COVID19 symptoms or vice versa because of overlapping molecular connections. We observed that the degree distribution of DDAN does not follow the scale-free property (Fig.2e). To find the exact topological nature, we measured network transitivity (*T_DDAN_* =0.4264) and average path length (*L_DDAN_ =2.0585*) of DDAN. These topological parameters were compared with the equivalent 1000 Erdős−Rényi random graphs. Our results show an average path length of DDAN is significantly less (p-value<0.0001), whereas transitivity, is significantly high (p-value<0.0001) compared to random graphs (*L_random_* = 2.44680 and *T_random_* = 0.0668) (Fig.2g). Further, we calculated small-worldness scalar (*S*) for DDAN as follows

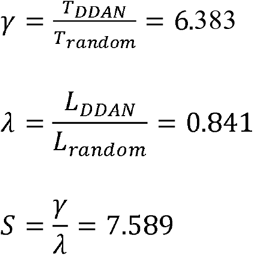

**Fig. 2.**
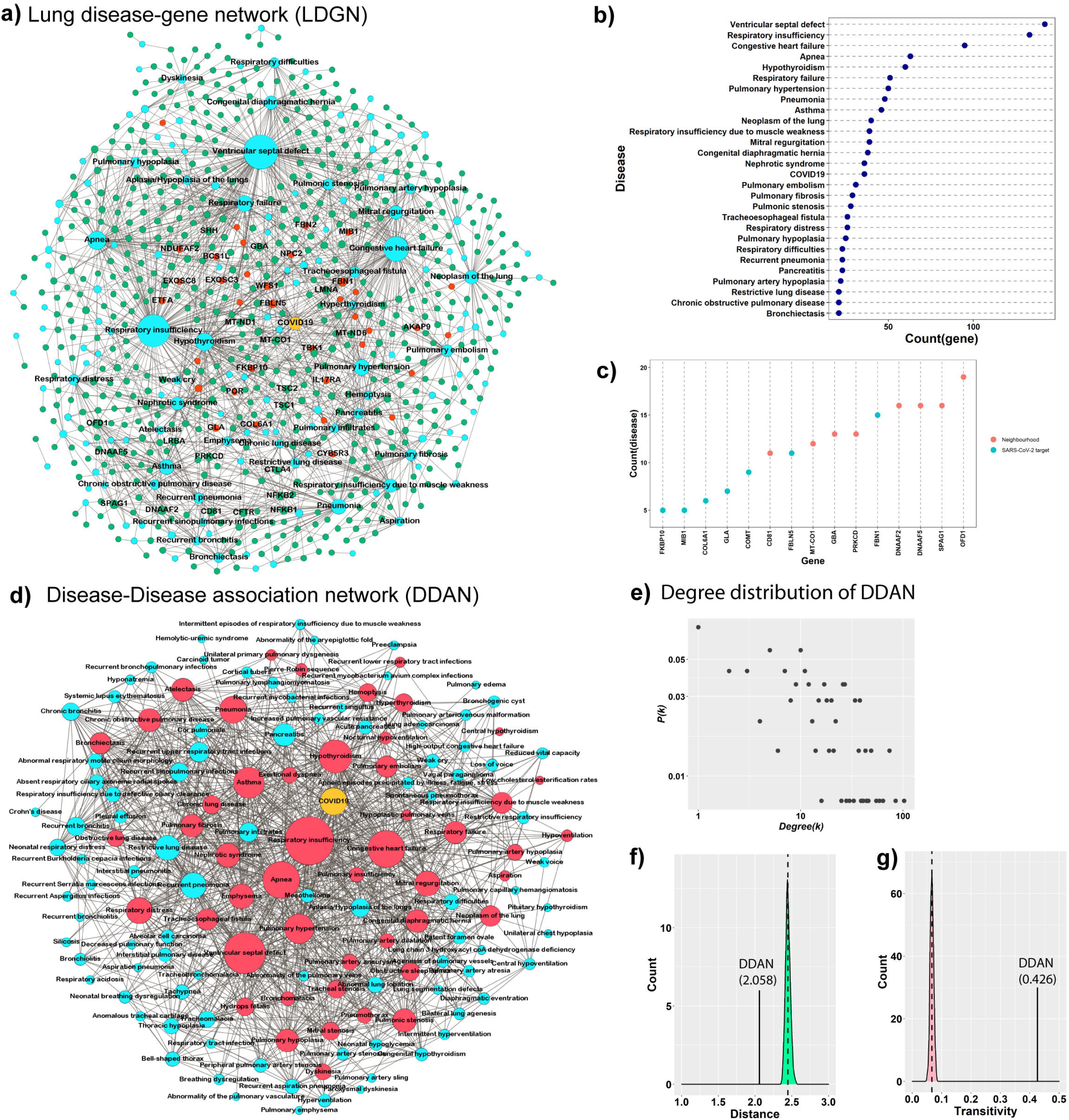
Disease-gene and disease-disease association network. a) Lung disease –gene network (LDGN), including COVID19 (yellow node). The network shows the SARS-CoV-2 targets (red) and neighborhood genes (green). b) & c) Dot plot shows the highly connected diseases (*k*>20) and genes in LDGN. d) Disease-disease association network(DDAN), red nodes represent the diseases that are directly direct linked to COVID19. e) Scatter plot shows the degree distribution of DDAN does not possess the scale-free property. f) The average path length between the diseases in DDNA and distribution of average path length of 1000 random networks (green). g) Transitivity of DDNA and distribution of transitivity of 1000 random networks (pink).

**Fig. 3.**
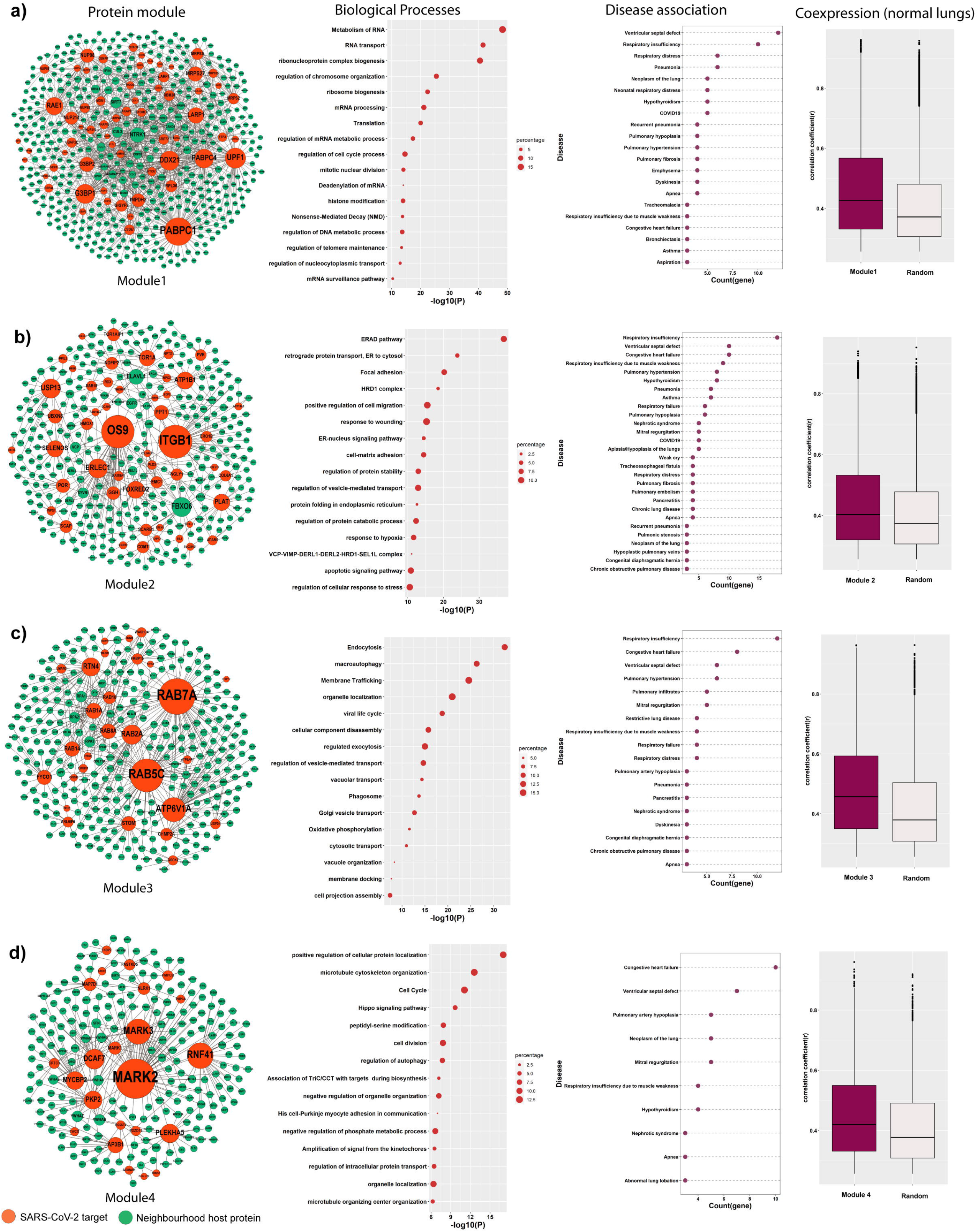
Community detection in STN and functional protein module. a,b,c &d show the modules 1 to 4, pathway and process enrichment analysis of each module, their disease associations, and coexpression of genes in each module in healthy lung tissue.

A network is said to be a small-world network if *S*>1[13]. Hence the topology of DDAN represents a small-world property, indicating any two diseases in DDAN have a high tendency to be interconnected and resulting in the overlapping pathogenesis between the diseases in DDAN. The molecular similarities between these lung disorders create a high-density comorbidity cluster and contribute to higher mortality in COVID19 patients. Therefore, it is necessary and a challenge to develop effective drugs to control the patient-specific risks of comorbidity in SARS-CoV-2 infection. However, it is difficult to select and prioritize the targets for treatment due to the several overlapping molecular connections. Therefore, we propose to target host functional protein modules associated with different disorders and hijacked by SARS-CoV-2.

### Functional protein modules preferentially hijacked by SARS-CoV-2 are linked to a broad range of lung disorders

Modularity in the network refers to the pattern of connectedness in which nodes are grouped into highly connected subsets [14]. One of the key features in the protein interaction network is that the tightly connected proteins within a community are mostly involved in similar biological functions [15]. Similarly, genes involved in related diseases are shown to be highly connected; moreover, diseases linked to common genes resulting in the formation of disease modules and comorbidity [16]. We have compared various community detection algorithms, i.e., fast-greedy, walktrap, louvain, leading eigenvector, and spinglass, to identify protein modules in STN [17,18]. Spinglass showed good partitioning, i.e., higher modularity score compared to other algorithms (see methods and supplementary table 4). Our findings are in agreement with previous studies by Rahiminejad et al. [19], where authors observed good partitioning of the functional protein module using spinglass in eukaryotes. Out of 21 modules, the top four protein modules were selected based on the presence of a large number of SARS-CoV-2 targets (>20) and gene ontology semantic similarity score (>0.2) of biological processes (supplementary table 5). A large number of the viral targets were considered because those modules are largely hijacked and strongly perturbed upon infection compared to other functional modules in the network. The modules were named as modules 1, 2, 3, and 4, and each module contains 63, 50, 28, 23 SARS-CoV-2 target protein**s**, respectively (Fig3). The biological process and pathway enrichment analysis show that module1, one the largest module, is mostly enriched with RNA metabolism, including transcription, mRNA processing, transport, mRNA deadenylation, and surveillance. Presumably, biological processes linked to module1 are hijacked by SARS-CoV-2 in the early stage of infection for the production of its RNA. Notably, the components of module1 are linked to 64 disorders, among which the highly connected are respiratory insufficiency, ventricular septal defect, respiratory distress, pneumonia, and neoplasm of the lung (Fig3a, 3rd column, supplementary table 6). It is worth noting that most of the diseases associated with module1are directly connected to COVID19 (Fig.2d and Fig.S2). On the other hand, hijacking module2 can predominantly affect the protein degradation (ERAD pathway, HRD1 complex, regulation of protein catabolic process), transport, folding and stability (retrograde protein transport, regulation of protein stability, VCP-VIMP-DERL1-DERL2-HRD1-SEL1L complex, regulation of intracellular transport, regulation of vesicle-mediated transport, protein folding in the endoplasmic reticulum). Module3 and module4 involve several processes, majorly cellular transport, localization, organization, and cell cycle. Modules 2, 3, and 4 are linked to a total of 79, 60, and 32 different disorders, respectively (supplementary table 6). Many clinical conditions such as ventricular septal defect, respiratory-related problems, neoplasm of the lung, apneas are associated with all modules, indicating a higher risk of the severe illness of patients on the onset of COVID19 infection. Besides, we found a wide spectrum of disorders of various classes such as neoplasm, neurological, and digestive system are associated with these modules (Fig S3). Gysi et al.[20] predicted the manifestation of SARS-CoV-2 in different human tissues could cause various disorders. Therefore not only lung-related disorders but comorbidity in various organs can also be a potential threat for COVID19 patients. To strengthen this observation, the pattern of coexpression of genes in functional modules was analyzed. Genes in the same functional module often show a high coexpression profile; therefore, we have calculated Pearson correlation coefficients of pairs of genes using gene expression data of healthy lung tissue from TCGA. The median value of the positive correlation between the genes in all modules is significantly higher (p-value< 0.0001) compared with the random gene set (Fig3, fourth column). Therefore theses modules can be identified as coexpress modules that share core transcriptional programs in the lung, indicating that their perturbation can result in a similar disease phenotype.

### Targeting the functional modules as a treatment strategy

We propose to target functional protein modules, hijacked by SARS-CoV-2, by drug repositioning. There are two main reasons to target these modules. Firstly, the binding of a drug to its target in a module will prevent the replication of the virus. Secondly, as a module is linked to several lung diseases, targeting a module can improve the severity of comorbidity. We identified 56 approved targets in the functional modules (red color nodes in Fig.4) from DrugBank [21]. Considering the complexity of COVID19, we also suggest using combination therapy to target multiple highly connected nodes simultaneously in the same or different functional modules (indicated by the arrow in Fig.4). For example, NTRK1 (*k*=43), and IMPDH2 (*k*=37) in module1, as well as PLAT (*k*=17), and COMT (*k*=10) in module2. Targeting these nodes can efficiently hinder the viral possession of these modules by rewiring the cellular network and can effectively reduce the growth of the virus. The target proteins suggested here, do not directly interact with SARS-CoV-2; rather, they are neighborhood nodes, as they are present in the same functional module, indicating their aggregation in the same network vicinity. Therefore binding of drugs to theses target can efficiently perturb the network modules as well as viral growth [22]. Importantly, the proposed targets should be tested and validated through clinical trials.

**Fig. 4.**
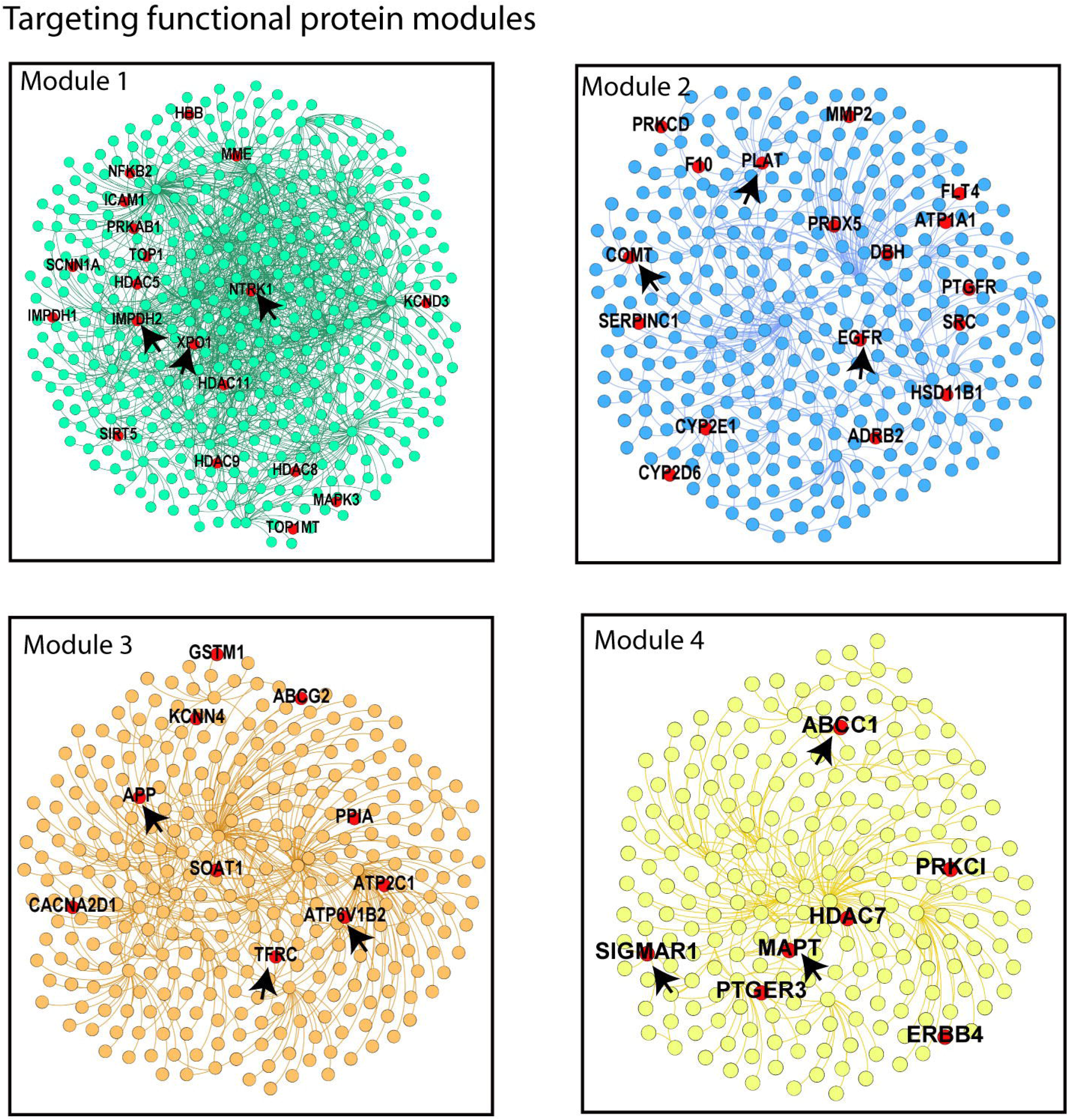
Targeting functional protein modules hijacked by SARS-CoV-2: The red nodes in each module indicate the FDA-approved targets, and an arrow indicates the highly connected node.

## Discussion

Currently, there is an urgent need for a speedy drug discovery or vaccine development to stop the infection and rapid transmission of SARS-CoV-2. Most alarming is that the aged COVID19 patients with comorbidity are in severe health risks worldwide. The present study has shown the risk of SARS-CoV-2 infection on the onset of various lungs related disorders and molecular basis of comorbidity by applying the principle of network biology. COVID19 appears to be a complex disease like cancer because of wide-ranging SARS-CoV-2 targets in the host cell, establishing the molecular connection with various lung-related disorders. The disease-gene and disease-disease association map, including COVID19, have shown overlapping molecular connections and high clustering of diseases in the same network vicinity, indicating a close pathobiological similarity. Indeed, the formation of a small-world network among lung diseases and COVID19 demonstrates a high-density comorbidity cluster. The observations strengthen our understanding of the molecular basis of the severe illness of COVID19 patients with comorbidity. It is now a great challenge to find the specific targets and potential drugs for patients with pre-clinical conditions because of large-scale molecular similarities of COVID19 with the other lung disorders. Therefore we suggest targeting the host functional protein modules, which are the origin of many lung disorders and primarily hijacked by the SARS-CoV-2. Perturbing these modules by repurposing FDA-approved drugs may rescue the host cellular machinery utilized by the virus for its replication. Realizing the complexity of SARS-CoV-2 infection, we further suggest testing multiple drugs or drug targeting various proteins, to improve clinical outcomes. Besides, patient-specific high-throughput transcriptomics data, in vitro, or in vivo assays are essential to establish the proper treatment strategy.

## Methods

### Construction of lung-specific PPI network of SARS-CoV-2 targets

Human lung tissue-specific interactome data was retrieved from the TissueNet v.2 database. To generate the tissue-specific PPIs, TissueNet v.2 synergizes between large-scale data of human PPIs and tissue-specific expression profiles. PPIs from four major PPI databases, BioGrid, IntAct, MINT and DIP, were obtained and consolidated. Then it integrated resulting PPIs with RNA-sequencing profiles of Genotype-Tissue Expression consortium (GTEx). We downloaded 168296 lung-specific interactions from TissueNet v.2 to construct SARS-CoV-2 targets interactome. Next, we obtained the list of 332 human proteins targeted by SARS-CoV-2 [8] and built a subnetwork, called the SARS-CoV-2 target network (STN). The 9 SARS-CoV-2 targets (AATF, CEP43, CISD3, MTARC1, NUP62, SRP19, THTPA, TIMM10B, TRIM59) do not have any interaction in the lung.

### Construction of lung-specific disease-gene and disease-disease network

The disease-gene association data in the lungs or effecting lungs were retrieved from the Gene ORGANizer (geneorganizer.huji.ac.il) [12]. Gene ORGANizer is a phenotype-based curated database that links human genes to the body parts they affect. Phenotypes that are classified by Human Phenotype Ontology (HPO) were considered with certain modifications. Disease-gene pairs that are not included but matching with the HPO phenotype were manually included. Aspirin-induced asthma and asthma were considered as asthma. Pulmonary emphysema, sarcoidosis and silicosis, and their associated genes were also added to the list. Finally, 6040 disease-gene pairs, which include a total of 184 various lung diseases, was mapped to STN. 618 out of 5050 nodes of STN were linked to 145 lung diseases, and 36 out of 618 genes were the direct target of SARS-CoV-2. These 36 genes were connected to COVID19 as new diseases-genes pair. Finally, a lung disease-gene association network, consisting of 1815 disease-gene pairs, including COVID19, was constructed. Disease-disease association network was derived from the lung disease-gene association network; two diseases were connected if they share one common gene. disgenet2r package [23] was used to study the association of disease classes with the functional protein module.

### Community detection

We applied fast-greedy, walktrap, louvain, leading eigenvector and spinglass on STN as an undirected, unweighted network. Theses community detection algorithms segregate the nodes into higher-density modules. Each of these algorithms optimizes an objective function i.e, modularity. Communities separated by spinglass were selected for subsequent analysis based on the modularity score and community size. Spinglass uses a random number generator to find communities. Therefore we ran Spinglass 10 times with different seed values. We compared the rand statistics between each run, and it showed the community structures are highly similar (>0.7) to each other [19,24].

### Process and pathway enrichment analysis and gene ontology (GO) Semantic similarity

Pathway and process enrichment analysis were performed using the Metascape [25]. GO Biological Processes, KEGG Pathway, and Reactome were used as ontology sources. GO semantic similarity between genes was measured by Wang *et al*.[26] method using GOSemSim package in R.

### Correlation analysis

TCGA gene expression datasets of human lung healthy tissues were downloaded from the UCSC Xena project (https://xenabrowser.net/datapages/) [27]. log_2_(RPKM +1) (RPKM: Reads Per Kilobase Million) transformed data of adjacent healthy tissue of 59 lung adenocarcinoma patients were retrieved, and Pearson correlation coefficient was computed to measure the coexpression levels using Hmisc Package in R.

### Computation of topological parameters

Average path length, transitivity, dyadicity, and Jaccard similarity coefficient were measured using igraph package in R. Average path length refers to the average length of pairwise shortest paths from a set of nodes to another set of nodes and transitivity (T) indicates the relative number of triangles in the graph, compared to a total number of connected triples of nodes. Dyadicity (D) measures the number of same label edges divided by the expected number of same label edges, and D> 1 indicates higher connectedness between the nodes with the same label. Jaccard similarity coefficient of two nodes is the number of common neighbors divided by the number of nodes that are neighbors of at least one of the two nodes being considered. Random network models were generated using the 1000 Erdös–Rényi random graph model of the same density. The random networks were compared with the original network by measuring the Z-score and p-value.

### Tools for data analysis and plotting

R packages tidyverse and stringr were used for data analysis, and plotting of graphs was done by ggplot2. Networks were visualized using Gephi. All statistical tests were performed using R.

## Acknowledgments

I thank Dr. Urmila Saxena (National Institute of Technology Warangal, Telangana) for critically reading the manuscript and her constructive comments and Dr. Subir Bhattacharjee (Purulia Government Medical College & Hospital, West Bengal) for discussion on lung disease. I also thank the National Institute of Technology Warangal for providing facilities.

## Supplementary information Supplementary Figures

**Fig.S1:** Number of genes linked to a lung disorder in LDGN

**Fig.S2:** Lung diseases directly connected to COVID19

**Fig.S3:** Various disease classes associated with functional protein modules

## Supplementary Tables

**Supplementary Table1:** Edge list of SARS-CoV-2 target network (STN) from TissueNet v.2 database

**Supplementary Table2:** Disease-gene association data of lungs from the ORGANizer database

**Supplementary Table3:** Lung disease-gene association data including COVID

**Supplementary Table4:** Comparison of different community detection algorithms applied to STN

**Supplementary Table5:** Protein modules generated using Spinglass algorithm

**Supplementary Table6:** Disease association with functional protein modules

## Reference

[1] Guan, W.J. et al. (2020). Clinical Characteristics of Coronavirus Disease 2019 in China. N Engl J Med 382, 1708–1720

[2] Mao, R., Liang, J., Shen, J., Ghosh, S., Zhu, L.R., Yang, H., Wu, K.C. and Chen, M.H. (2020). Implications of COVID-19 for patients with pre-existing digestive diseases. Lancet Gastroenterol Hepatol 5, 426–428.

[3] Li, B., Yang, J., Zhao, F., Zhi, L., Wang, X., Liu, L., Bi, Z. and Zhao, Y. (2020). Prevalence and impact of cardiovascular metabolic diseases on COVID-19 in China. Clin Res Cardiol 109, 531–538.

[4] Guan, W.J. et al. (2020). Comorbidity and its impact on 1590 patients with Covid-19 in China: A Nationwide Analysis. Eur Respir J. DOI: 10.1183/13993003.00547-2020

[5] Das, A.B. (2019). Disease association of human tumor suppressor genes. Mol Genet Genomics 294, 931–940.

[6] Goh, K.I., Cusick, M.E., Valle, D., Childs, B., Vidal, M. and Barabasi, A.L. (2007). The human disease network. Proc Natl Acad Sci U S A 104, 8685–90.

[7] Zheng, C. and Xu, R. (2018). Large-scale mining disease comorbidity relationships from post-market drug adverse events surveillance data. BMC Bioinformatics 19, 500.

[8] Gordon, D.E. (2020). A SARS-CoV-2 protein interaction map reveals targets for drug repurposing. Nature, DOI: 10.1038/s41586-020-2286-9.

[9] Pachetti, M. et al. (2020). Emerging SARS-CoV-2 mutation hot spots include a novel RNA-dependent-RNA polymerase variant. J Transl Med 18, 179.

[10] Basha, O., Barshir, R., Sharon, M., Lerman, E., Kirson, B.F., Hekselman, I. and Yeger-Lotem, E. (2017). The TissueNet v.2 database: A quantitative view of protein-protein interactions across human tissues. Nucleic Acids Res 45, D427–D431.

[11] Wang, Q. et al. (2012). Community of protein complexes impacts disease association. Eur J Hum Genet 20, 1162–7.

[12] Gokhman, D., Kelman, G., Amartely, A., Gershon, G., Tsur, S. and Carmel, L. (2017). Gene ORGANizer: linking genes to the organs they affect. Nucleic Acids Res 45, W138–W145.

[13] Bassett, D.S. and Bullmore, E.T. (2017). Small-World Brain Networks Revisited. Neuroscientist 23, 499–516.

[14] Wagner, G.P., Pavlicev, M. and Cheverud, J.M. (2007). The road to modularity. Nat Rev Genet 8, 921–31.

[15] Tripathi, S., Moutari, S., Dehmer, M. and Emmert-Streib, F. (2016). Comparison of module detection algorithms in protein networks and investigation of the biological meaning of predicted modules. BMC Bioinformatics 17, 129.

[16] Barabasi, A.L., Gulbahce, N. and Loscalzo, J. (2011). Network medicine: a network-based approach to human disease. Nat Rev Genet 12, 56–68.

[17] Reichardt, J. and Bornholdt, S. (2006). Statistical mechanics of community detection. Phys Rev E Stat Nonlin Soft Matter Phys 74, 016110.

[18] Blondel, V.D., Guillaume, J.-L., Lambiotte, R. and Lefebvre, E. (2008). Fast unfolding of communities in large networks. Journal of Statistical Mechanics 2008, P10008.

[19] Rahiminejad, S., Maurya, M.R. and Subramaniam, S. (2019). Topological and functional comparison of community detection algorithms in biological networks. BMC Bioinformatics 20, 212.

[20] Gysi, M.D. et al. (2020). Network Medicine Framework for Identifying Drug Repurposing Opportunities for COVID-19. arXiv:2004.07229

[21] Wishart, D.S. et al. (2018). DrugBank 5.0: a major update to the DrugBank database for 2018. Nucleic Acids Res 46, D1074–D1082.

[22] Yildirim, M.A., Goh, K.I., Cusick, M.E., Barabasi, A.L. and Vidal, M. (2007). Drug-target network. Nat Biotechnol 25, 1119–26.

[23] Pinero, J. et al. (2017). DisGeNET: a comprehensive platform integrating information on human disease-associated genes and variants. Nucleic Acids Res 45, D833–D839.

[24] Reichardt, J. and Bornholdt, S. (2006). Statistical mechanics of community detection. Phys Rev E Stat Nonlinear Soft Matter Phys 74, 016110.

[25] Zhou, Y., Zhou, B., Pache, L., Chang, M., Khodabakhshi, A.H., Tanaseichuk, O., Benner, C. and Chanda, S.K. (2019). Metascape provides a biologist-oriented resource for the analysis of systems-level datasets. Nat Commun 10, 1523.

[26] Yu, G., Li, F., Qin, Y., Bo, X., Wu, Y. and Wang, S. (2010). GOSemSim: an R package for measuring semantic similarity among GO terms and gene products. Bioinformatics 26, 976–8.

[27] Goldman, M. et al. (2019). The UCSC Xena platform for public and private cancer genomics data visualization and interpretation. bioRxiv. DOI: https://doi.org/10.1101/326470.

