## Supplementary Figures for "Lung Disease Network Reveals the Impact of Comorbidity on SARS-CoV-2 infection"

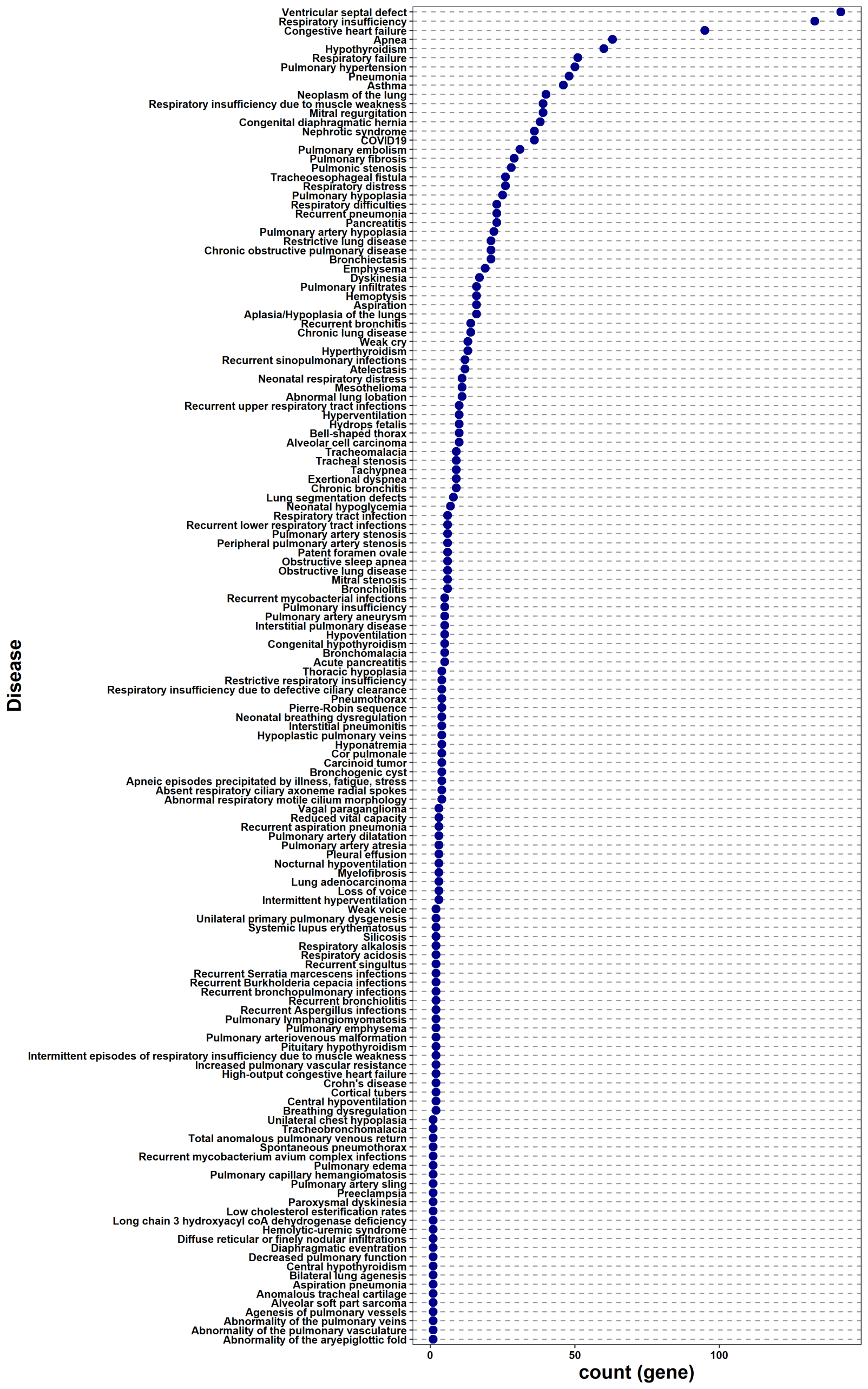

**Fig. S1:** Dot plot shows the number of genes associated with a lung disorder in LDGN.

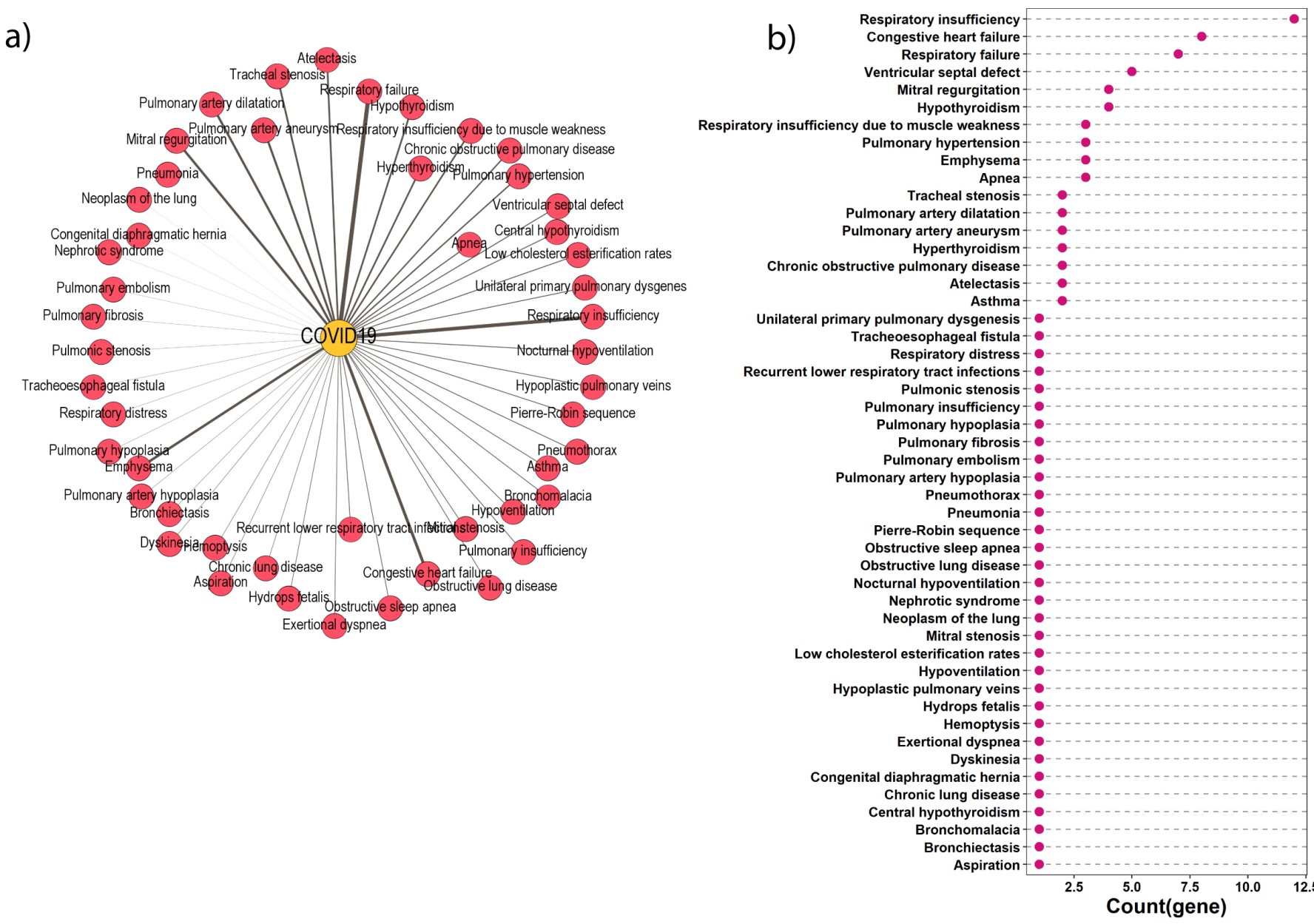

**Fig.S2** a) Network shows the lung disorders which are directly connected to COVID19. The thickness of the edge in the network is proportional to the Jaccard similarity coefficient. b) Dot plot shows the number of shared genes between COVID other lung disorders.

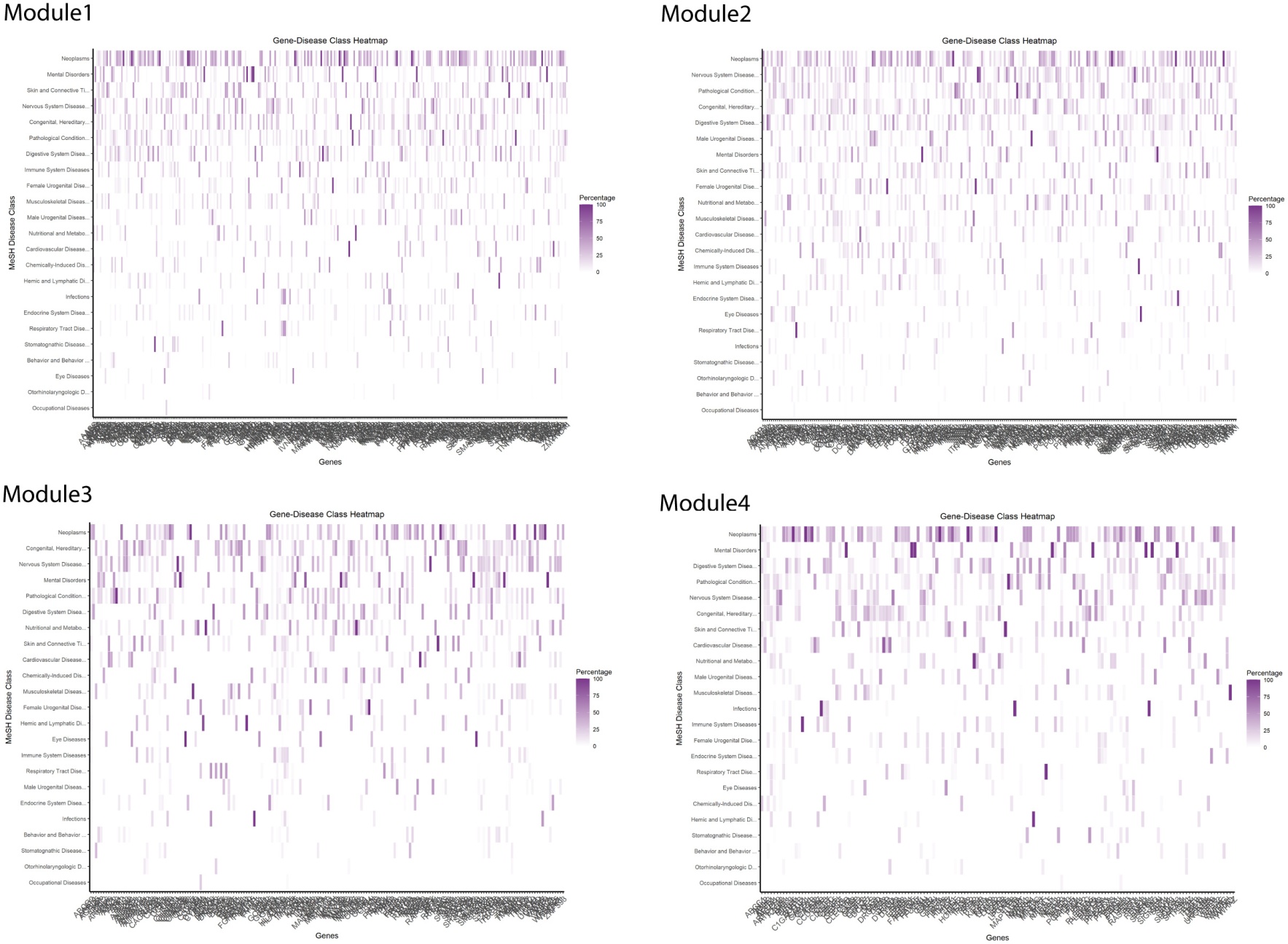

**Fig.S3** Heat map shows functional protein modules are associated with different disease classes.
